# Reverse Correlation Characterizes More Complete Tinnitus Spectra in Patients

**DOI:** 10.1101/2023.10.06.561243

**Authors:** Nelson V. Barnett, Alec Hoyland, Divya A. Chari, Benjamin Parrell, Adam C. Lammert

**Author notes:** The authors declare no competing interests.

## Abstract

**Goal:** We validate a recent reverse correlation approach to tinnitus characterization by applying it to individuals with clinically-diagnosed tinnitus.

**Methods:** Two tinnitus patients assessed the subjective similarity of their non-tonal tinnitus percepts and random auditory stimuli. Regression of the responses onto the stimuli yielded reconstructions which were evaluated qualitatively by playing back resynthesized waveforms to the subjects and quantitatively by response prediction analysis.

**Results:** Subject 1 preferred their resynthesis to white noise; subject 2 did not. Response prediction balanced accuracies were significantly higher than chance across subjects: subject 1: 0.5963, subject 2: 0.6922.

**Conclusion:** Reverse correlation can provide the foundation for reconstructing accurate representations of complex, non-tonal tinnitus in clinically diagnosed subjects. Further refinements may yield highly similar waveforms to individualized tinnitus percepts.

**Impact Statement:** Characterization of tinnitus sounds can help clarify the heterogeneous nature of the condition and link etiology to subtypes and treatments.

## I. Introduction

**T**innitus, the internal presence of an externally nonexistent sound, is a pervasive condition [1], [2] that varies greatly in perception, duration, severity, and etiology [3], [4]. Understanding heterogeneity of tinnitus perception is a longstanding goal in tinnitus research that may open the door to establishing subtypes and endotypes, improving treatment options and outcomes [5]–[8], and identifying specific behavioral and neurological mechanisms [9]–[11]. Accurate characterization of individualized tinnitus sounds enables a better understanding of this heterogeneity [12]–[14].

*Reverse correlation* (RC) is a widely established method for estimating internal perceptual representations using random stimuli and subjective responses [15]–[17]. Recent work has shown that RC works well for reconstructing the frequency spectra of tinnitus-like percepts in healthy controls [18]. Here, we assess the viability of RC for reconstructing tinnitus spectra in clinically-diagnosed tinnitus subjects. Two tinnitus patients completed an RC experiment, after which we conducted reconstruction validation via response prediction analysis [19] and subjective resynthesis assessments. This work demonstrates the effectiveness of RC for estimating tinnitus percept spectra in clinical subjects.

## II. Materials and Methods

### A. Subjects

Two subjects with clinically diagnosed tinnitus were recruited through the UMass Audiology clinic and referred to WPI for this study. Subjects completed a questionnaire regarding the frequency, severity, and characteristics of their tinnitus. Both subjects rated their tinnitus severity as 5 on a scale from 1 (most mild) to 10 (most severe). Subject 1 reported their tinnitus as “ringing, roaring, and humming.” Subject 2 reported hearing “static, hissing, ocean waves, and whooshing.” Both subjects had pure-tone hearing thresholds *<* 60 dB HL between 250 and 2000 Hz.

### B. Stimuli

For each stimulus, the frequency space between [100, 13000] Hz was divided into *b* = 32 Mel-spaced bins. [6, 16] bins were randomly assigned 0 dB, the rest *−*100 dB. All frequencies were assigned random phase, after which inverse Fourier transform of the spectrum yielded a 500 ms stimulus waveform.^1^

**TABLE 1.** Balanced accuracy values for 5-fold cross-validation response prediction analysis. RC is comparable to both LWLR and XGBoost for subject 1 and outperforms both for subject 2. Values generally increase with % “yes” responses, with the exception of XGBoost. Balanced accuracies on the training data are consistently greater than testing but less than 1, indicating minimal overfitting.

| Subject | % “yes” | Training |  |  | Testing |  |  |
| --- | --- | --- | --- | --- | --- | --- | --- |
|  |  | RC | LWLR | XGBoost | RC | LWLR | XGBoost |
| 1 | 13.000 | 0.6641 | 0.6961 | 0.8763 | 0.5963 | 0.6080 | 0.6429 |
| 2 | 39.625 | 0.7156 | 0.7646 | 0.7607 | 0.6922 | 0.6752 | 0.6417 |

### C. Experiment

Subjects completed the experiment in an IAC 120-A double wall audiometric booth, listening over Sennheiser HD 560S headphones (Sennheiser electronic GmbH & Co. KG, Wedermark, Germany) at a self-determined comfortable level. Presentation level was not recorded. Subjects were instructed to answer “yes” to stimuli that either “completely sounded like” or “had elements that resembled” their tinnitus (*cf*. Fig. 1). Both subjects completed 10 blocks of 80 trials (*p* = 800) with self-controlled breaks between each block. Procedures were approved by the UMass IRB.

**Fig. 1.**
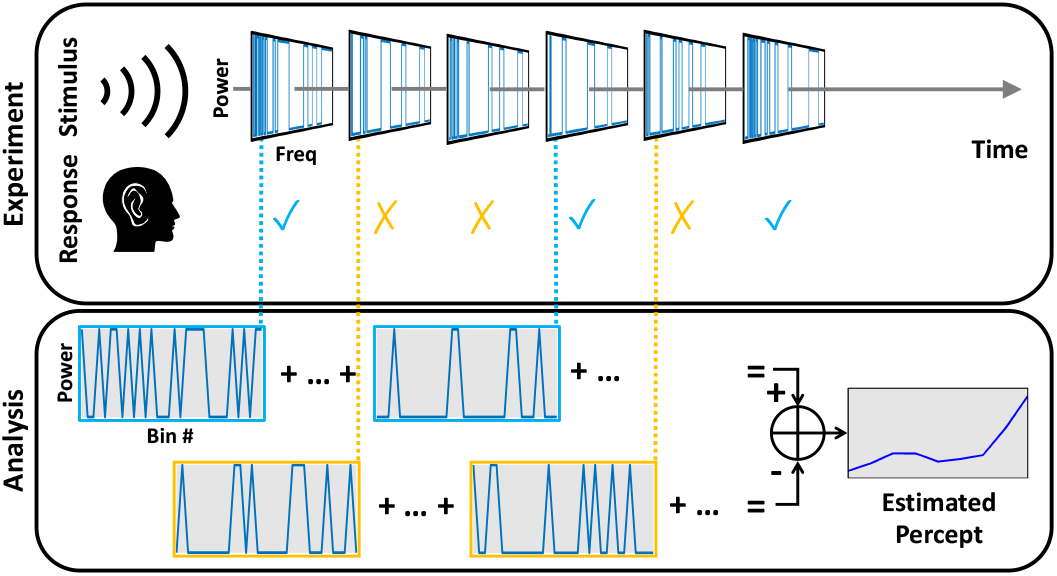
Diagram of the experimental protocol. Subjects listen to a series of random stimuli and respond either “yes” or “no” depending the perceived similarity to their tinnitus. The recorded stimulus-response pairs are used to form an estimate of the subject’s tinnitus percept.

### D. Reconstruction

RC was implemented to reconstruct the internal tinnitus representation given a stimulus matrix Ψ *∈*ℝ^*p×b*^ and a response vector *y ∈*{*−*1, 1}^*p*^, where 1 represents a “yes”, *−*1 a “no”, *p* is the number of trials, and *b* is the number of bins. RC assumes the subject response model:

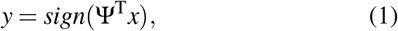

where *x ∈*ℝ^*b*^is the internal representation of interest. Assuming uncorrelated stimulus dimensions, inverting this model yields a restricted version of the Normal equation [16]:

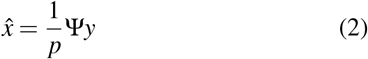

### E. Resynthesis

500 ms resynthesis waveforms were generated from binned reconstructions (Eq. 2) by rescaling the reconstructions to a 20 dB range, assigning random phase, and applying an inverse Fourier transform.

### F. Qualitative Assessment

Subjects were played unlabeled, repeatable sounds of their resynthesis and white noise in a randomized order once, ranking the statement “This sounds like my tinnitus” on a scale from 1 (strongly disagree) to 5 (strongly agree) after each.

### G. Prediction

An established quantitative measure of reconstruction quality is the accuracy of subject response predictions on the basis of those reconstructions [19]. To calculate this, stimuli-response pairs of experiment data were partitioned according to a 5-fold cross-validation scheme. For each fold, a reconstruction, 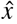, was made by applying Eq. 2 to Ψ_*train*_ and *y*_*train*_. Predicted responses, 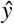, were generated via Eq. 1, where 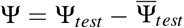, 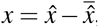, and 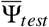 is the mean of Ψ_*test*_ in the second dimension.

Balanced accuracy (Eq. 3) was used as the primary validation metric. Robustness to class imbalance is critical because subjects responded “yes” less often (see Table I).

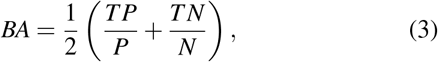

where *TP* and *TN* are the number of true positive and negative predictions, and *P* and *N* are the number of positive and negative responses.

In the absence of repeated exposures to the same stimuli, it can be challenging to establish an upper bound on prediction accuracy [19]. We therefore contextualize response prediction accuracies by presenting accuracies on the training data, an indication of performance with overfitting, and prediction results from two nonlinear models—locally weighted linear regression (LWLR) and XGBoost [20]—which may indicate an upper bound on performance.

The same cross-validation scheme was run for LWLR. The diagonal weight matrix, *W*, was found via:

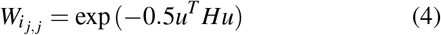

for *i ∈* 1, …, *p*_*test*_ and *j ∈*1, …, *p*_*train*_, where *H* = *hI*_*b*_ for *h ∈*ℝ and 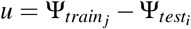. Local regression coefficients were determined using Eq. 5 with ε = 0.01 and estimations 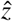 (Eq. 6) were converted into response predictions via Eq. 7.

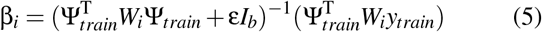

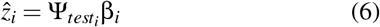

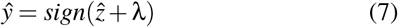

Both *h* and λ were optimized each fold using a development set.

XGBoost using regression, squared error loss, and a gradient boosted tree method was run in Julia over the same cross-validation protocol. Estimations were converted to response predictions using Eq. 7 with λ, number of boost rounds, and both *l*_1_ and *l*_2_ regularizers similarly optimized on a development set.

## III. Results

Prediction BAs using RC for both subjects were well above 0.5, with RC outperforming both LWLR and XGBoost for subject 2 (Table I). Both RC and LWLR results improve with percent “yes” responses, while XGBoost performs consistently between subjects. Distinct differences are observable between the subjects’ reconstructions (Fig. 2), indicating subjects were attuned to different elements of the stimuli. Subject 1 rated their resynthesis a 4 and white noise a 2. Subject 2 rated their resynthesis a 1 and white noise a 3 (see Section II-F).

**Fig. 2.**
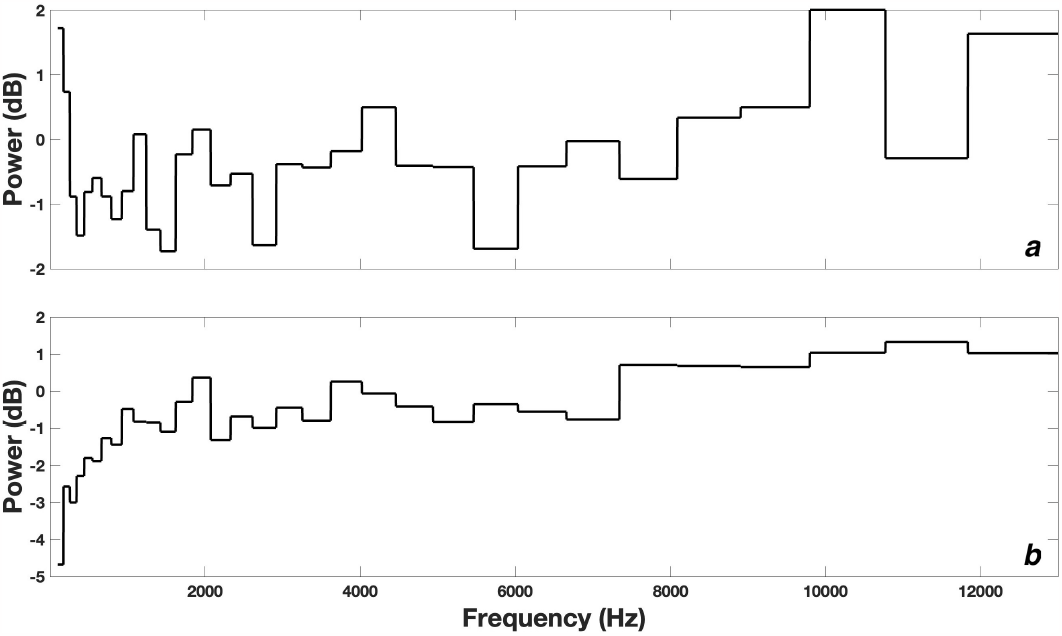
Reconstructions for *(a)* subject 1 and *(b)* subject 2 generated via Eq. 2.

## IV. Conclusion

This study demonstrates the effectiveness of RC in reconstructing accurate tinnitus spectra from individuals suffering from tinnitus. Reconstruction accuracy is well supported by response prediction analysis (Table I), for which RC performs close to an upper bound estimated by two nonlinear models. Furthermore, BA on training data shows minimal overfitting. Altogether, this indicates that the majority of the performance loss is likely due to irreducible error in the subjects’ responses. The reconstructed tinnitus spectra in Figure 2 therefore represent the first of their kind for individuals experiencing non-tonal, spectrally complex tinnitus.

The heterogeneity of tinnitus implies that two subjects cannot adequately represent the range of tinnitus experiences. In a larger sample population, it will be critical to minimize subject response error to reliably reconstruct accurate representations of complex tinnitus spectra. Methodological refinement to the stimuli may significantly improve this. While subjective resynthesis evaluations were mixed, high response prediction BAs point to a sound experimental protocol and reconstruction method. Improvements in the resynthesis technique are likely to greatly boost subjective assessment scores.

## Footnotes

1 Software for the experiments and analysis was written in MATLAB and is freely available at https://github.com/The-Lammert-Lab/tinnitus-reconstruction

